# Comparative Transcriptome Profiling Provides Insights into the Growth Promotion Activity of *Pseudomonas fluorescens* strain SLU99 in Tomato and Potato Plants

**DOI:** 10.1101/2023.01.08.523145

**Authors:** Nurul Atilia Shafienaz binti Hanifah, Farideh Ghadamgahi, Samrat Ghosh, Rodomiro Ortiz, Stephen C. Whisson, Ramesh R. Vetukuri, Pruthvi B. Kalyandurg

## Abstract

The use of biocontrol agents with plant growth-promoting activity has emerged as an approach to support sustainable agriculture. During our field evaluation of potato plants treated with biocontrol rhizobacteria, four bacteria were associated with increased plant height. Using two important solanaceous crop plants, tomato and potato, we carried out a comparative analysis of the growth-promoting activity of the four bacterial strains: *Pseudomonas fluorescens* SLU99, *Serratia plymuthica* S412, *S. rubidaea* AV10, and *S. rubidaea* EV23. Greenhouse and *in vitro* experiments showed that *P. fluorescens* SLU99 promoted plant height, biomass accumulation, and yield of potato and tomato plants, while EV23 promoted growth in potato but not in tomato plants. SLU99 induced the expression of plant hormone-related genes in potato and tomato, especially those involved in maintaining homeostasis of auxin, cytokinin, gibberellic acid and ethylene. Our results reveal potential mechanisms underlying the growth promotion and biocontrol effects of these rhizobacteria and suggest which strains may be best deployed for sustainably improving crop yield.

## 1 Introduction

The plant rhizosphere is a significant carbon sink (Strand et al., 2008; Drigo et al., 2010), and represents an ecosystem hosting a diverse microbial community, including a variety of phytopathogens. The co-evolution of plants and microbial communities has led to the formation of complex symbiotic associations (Delaux and Schornack, 2021; Abdelfattah et al., 2022). Several studies have indicated that plants recruit specific microbial communities through root exudates, positively regulating the rhizosphere (Besserer et al., 2006; De Weert et al., 2007; Rudrappa et al., 2008). Through secreting secondary metabolites such as pyrrolnitrin and fusaricidin (Burkhead et al., 1994; Vater et al., 2017; Liu et al., 2020), outer membrane vesicles (McMillan et al., 2021) and phytohormones (Tsukanova et al., 2017), microbial communities assist plants in their defence against pathogens by priming plant defences while keeping severe immune responses to a minimum (Van Wees et al., 2008). Termed biological control agents (BCA), several bacterial genera including *Pseudomonas, Serratia, Bacillus*, and *Azospirillum* compete in the rhizosphere environment with pathogens through siderophore biosynthesis, antibiosis, lytic enzyme production and detoxification, thus protecting the plant (Loper, 1988; Loper and Henkels, 1999; Compant et al., 2005). Some species of rhizospheric BCA, known as plant growth-promoting rhizobacteria (PGPR), have long been known to enhance plant growth through the secretion of auxin (Přikryl et al., 1985), cytokinin (Tien et al., 1979; Timmusk et al., 1999), 1-aminocyclopropane-1-carboxylate (ACC) deaminase (Glick et al., 1994), nitrogen fixation (Bal and Chanway, 2012), volatile organic compound (VOC) production (Tahir et al., 2017), and phosphate solubilization (Son et al., 2006; Olanrewaju et al., 2017).

Many PGPR modulate root architecture, promote shoot elongation, and increase chlorophyll content and photosynthetic efficiency (Diagne et al., 2020). One of the most common alterations induced by PGPR in the root system architecture is the increased formation of lateral roots (LR) and root hairs (RH), thus assisting in improved water and nutrient acquisition from the soil. Changes in root system architecture (RSA) could be caused by PGPR interference with the signalling in the main hormonal pathways regulating plant root development, namely auxin and cytokinin (Ali et al., 2009; Dodd et al., 2010). PGPRs also promote plant growth by reducing stress-related ethylene levels via ACC deaminase activity (Glick et al., 1998; Hardoim et al., 2008; Amna et al., 2019). PGPR have been demonstrated to be beneficial for several important crop species including tomato, apple, grapes, and cereals such as rice, wheat and maize (Kumari et al., 2019).

*Pseudomonas* spp. are Gram-negative, chemoheterotrophic, motile rod-shaped bacteria that are adapted to a wide range of ecological niches. Extensive research has shown that several *Pseudomonas* strains function as PGPR through diverse metabolic capabilities. *P. putida* isolate WCS358 (Zamioudis et al., 2013) and *P. chlororaphis* subsp. *aurantiaca* strain JD37 (Fang et al., 2013) isolated from potato rhizosphere promote plant growth in *Arabidopsis thaliana* and maize, respectively. *P. putida* isolate GR12-2 enhances root elongation through the ethylene pathway (Glick et al., 1994) and root branching by producing cyclodipeptides that modulate auxin responses (Ortiz-Castro et al., 2020). Furthermore, we have recently demonstrated that *P. aeruginosa* encodes genes for tryptophan biosynthesis and indole-3-acetic acid (IAA) synthesis, and promotes tomato growth (Ghadamgahi et al., 2022). *P. fluorescens* strain WCS365 has been shown to colonize both tomato and potato rhizospheres (Mercado-Blanco and Bakker, 2007). *P. fluorescens* strain WCS417 inhibits primary root elongation while stimulating LR and RH development through auxin signalling (Zamioudis et al., 2013).

*Serratia* sp. are Gram-negative bacteria belonging to the family Enterobacteriaceae. Several species of *Serratia* are reported to enhance plant growth, such as *S. fonticola* strain AU-P3 and *Serratia* sp. SY5 that promote growth in pea and maize plants, respectively (Koo and Cho, 2009; Devi et al., 2013). Recently, it was reported that *Serratia rubidaea* strain ED1 promotes seed germination in quinoa (Mahdi et al., 2021).

A shift in societal acceptance away from using harmful and often expensive chemical fertilizers to a safer and natural alternative, combined with the increasing challenges of climate change and the growing world population, has resulted in PGPR drawing increased attention. The effects of PGPR-plant interactions on molecular mechanisms are complex and may vary depending on the strain or consortia of strains, plant species, and receiving environment (Tsukanova et al., 2017). Understanding how a beneficial bacterium improves the growth and health of a specific crop in a specific environment is therefore critical. The strains used in this study were isolated from potato and tomato rhizosphere soils. *In vitro* antagonistic analysis indicated a strong antagonistic activity against plant pathogens, including *P. colocasiae* (Kelbessa et al., 2022), *P. infestans* and *R. solani* (unpublished). The aim of this study was to conduct a holistic study on the effect of PGPR on potato and tomato growth and select the best-performing PGPR to gain insights into their mechanism of action. To that end, we carried out a field trial with six bacteria with biological control activity on potato plants in a high disease pressure *P. infestans* infested field. Selected strains associated with increased potato growth were then tested under greenhouse conditions for growth-promoting activity.

The complex effects of PGPR on plant gene expression, including genes that play key roles in signal transduction, metabolism, and catabolism of plant hormones have been reported in several studies (Shen et al., 2013; Tsukanova et al., 2017; Wei et al., 2020). However, studies on global transcriptomic changes induced by PGPR treatment in the context of plant growth and development are limited. Moreover, to our knowledge, there is no report on the transcriptome profiling of potato plants under PGPR treatment. Understanding how plants including potato and tomato respond to PGPR treatment will greatly empower the usage of PGPR towards enhancing sustainable agriculture. To that end, we carried out transcriptome profiling of potato and tomato roots and leaves treated with the bacterial supernatant to gain insight into the growth promotion mechanisms of *P. fluorescens*.

## 2 Materials and methods

### 2.1 Field trial

Field experiments were carried out in the summer of 2020 using the potato cultivar Kuras. A total of six bacterial isolates that displayed antagonistic activity against *Phytophthora infestans* under lab conditions were assessed for their plant growth promotion activity. These strains include *Pseudomonas fluorescens* SLU99, *Serratia rubidaea* EV23 and AV10, *S. plymuthica* S412 and AS13, and *S. proteamaculans* S4. One single colony of each bacterial strain was cultured in Luria-Bertani (LB) liquid for 16 h at 28°C. Then the cultures were centrifuged at 3000 x g for 10 min and the bacterial cells were resuspended in 5 L sterile water to a final OD_600_ 0.2. The bacterial suspension was transferred to a 5 liter pressure sprayer for field application, water treatment were used as a control. The testing site was Mosslunda, near the city of Kristianstad (55°58’00.3”N 14°07’03.0”E) where the average daily temperature ranged between 12 and 18°C and average monthly precipitation was 42-64 mm, with an average daylength was 17.5 hours. The experiment was carried out in a plot with four rows of 10 meters in width and 15 meters in length. The experimental design consisted of randomized block design with four blocks. Each block contained all eight treatments distributed randomly, with 10 plants per treatment. The potato plants were sprayed with six biocontrol bacteria (individual treatments) once every two weeks for a period of 8 weeks.

### 2.2 Greenhouse and PGPR treatment

To evaluate the effects of selected rhizobacteria on tomato and potato plant growth, a pot experiment was conducted for each crop in two separate plant cultivation chambers (biotron). Nine and ten biological replicates were used for tomato and potato, respectively. The tomato seeds were surface sterilized using 3% bleach followed by washing with sterile water three times before sowing in a 105-hole seed tray filled with nursery soil. At 25 days after sowing, uniform four-leaf stage seedlings were transplanted into 1 L pots containing 325 g of nursery soil (1 seedling/pot). For bacterial inoculations, strains SLU99, EV23, AV10 and S412 were cultured as described in the previous section. Thirty-five milliliters of bacterial suspensions at OD_600_ 0.2 prepared with 1 × phosphate-buffered saline (PBS) (Thermo Scientific, Waltham, MA, USA) or only 1 × PBS (control treatment) was added to the roots of tomato seedlings before they were covered with soil. For potato, all seed tubers were first cleaned with water, treated with 100 ppm gibberellic acid, and dried for eight days to promote bud initiation. Prior to planting in 2 L pots (containing 650 g nursery soil), the seed tubers were inoculated with bacteria suspension at OD_600_ 0.2 prepared with 1 × PBS or with only 1 × PBS for 15 min.

The inoculation was repeated twice, once at 10 and 30 days after transplanting (DAT) for tomato or days after planting (DAP) for potato with 35 ml bacterial suspension (OD_600_ 0.2) per plant in both crops to ensure the growth of bacteria in the soil. The liquid was poured 2-3 cm from each plant’s base and 5 cm deep in the soil. For the duration of the experiments, the plants were grown in two separate growth chambers in 16 h light/8 h dark photoperiods supplemented with 250 µmol m-2 s-1 light, temperatures of 20-23°C and 60% humidity for 60 days. The plant positions within the growth chambers were randomly rearranged at least twice a week.

### 2.3 Phenotyping

To determine the effects of selected rhizobacteria on plant growth, parameters of plant height, total plant dry weight and yield were measured. Plant height was measured from the soil surface to the tip of the plant every ten days from planting until harvest. The yield of tomato and potato were measured by total fruit or tuber number and total fresh weight. Firstly, fruit or tubers were harvested, counted, and weighed. Then, the shoot and root systems were separated. Leaves were separated from the plant by cutting the leaf petiole, and the stems were cut. The number of leaves was counted and recorded. The intact roots were gently shaken to remove soil, briefly washed with water, and gently blotted dry using paper towels. Then, immediately the roots, leaves, stems, and fruits/tubers were separately placed in paper bags and oven-dried at 65°C until they reached constant weight. Finally, the dry weight was measured using a precision scale and recorded.

### 2.4 Chlorophyll content index measurement

The chlorophyll content of tomato and potato plants grown in the growth chamber were measured at harvest using the MC-100 Chlorophyll Concentration Meter (Apogee Instruments Inc., Logan, UT, USA). The MC-100 was calibrated to measure chlorophyll concentration with units of chlorophyll content index (CCI) and zeroed before commencing measurement according to the manufacturer’s instructions (Apogee Instruments, 2018). The generic equation was used in the measurement of the relative chlorophyll content of both crops. For tomato plants, the measurements were made by clipping the sensor onto the second terminal leaflet on the fifth fully expanded leaf from the top of each plant (Matsuda et al., 2014). For potato plants, the measurements were made on the top point of the top leaflet of the 4th compound leaf (Li et al., 2012).

### 2.5 Soil sampling and analysis

To evaluate the effects of rhizobacteria applications on soil nutrient content, soils were collected and pooled from all pots of each bacterial treatment for analysis. For the control treatment, soils were collected from eight randomly non-inoculated plants. The soil samples were sent to the LMI AB testing laboratory (Helsingborg, Sweden) for analysis. The soils sampled before planting and at harvest were analyzed for pH, soil organic matter (SOM), electrical conductivity (EC), total nitrogen, available phosphorus (P), and available potassium (K).

### 2.6 *In vitro* growth conditions for plants

Tomato seeds or potato explants were grown in half-strength Murashige and Skoog (MS) culture medium (M0221; Duchefa Biochemie B.V, Haarlem, The Netherlands), pH 5.8, supplemented with 0.25% (w/v) Phytagel (Sigma-Aldrich Co., St. Louis, MO, United States), 0.05% (w/v) 2-(N-morpholino)ethanesulfonic acid (MES) and 1% sucrose (w/v) (Duchefa Biochemie B.V, Haarlem, The Netherlands). For SLU99 treatment, bacteria were cultured overnight in LB until an OD_600_ of 2 was reached. Cell-free supernatant was obtained by centrifuging at 3000 x g for 10 min, filtered through a 0.22-μm filter and mixed with ½ MS medium at 1:10 ratio before plating. Tomato var Moneymaker seeds were obtained from Plantagen AB and were surface sterilized with 3% (v/v) sodium hypochlorite for 5 min and washed three times with sterile water. Seeds were then plated either on control or SLU99 supernatant supplemented ½ MS plates and stratified for 48 h at 4°C. For potato plants, the explants from sterile potato plants were grown either on mock treated or SLU99 supernatant supplemented ½ MS media in plant tissue culture boxes. The plants were grown in a growth chamber with 16 h light/8 h dark photoperiods and temperatures of 20-22°C.

### 2.7 Transcriptome analysis

For RNA-seq analysis, leaf or root samples were collected for each treatment and frozen in liquid nitrogen. RNA was extracted using the RNeasy Mini Kit (Qiagen). The quality and integrity of the RNA was measured using Agilent Bioanalyzer 2100 (Agilent, California, USA). Library preparation was carried out using a TruSeq RNA poly-A selection kit (Illumina, Inc.). Sequencing was performed at National Genomics Infrastructure (NGI), Stockholm on an Illumina NovaSeq6000 S4 platform. Adapter sequences and poor quality reads (<Q30) were removed using BBduk (Bushnell et al., 2017). Next, cleaned data were fed into STAR (Dobin et al., 2013) for alignment against the reference genome. FeatureCounts (Liao et al., 2014) was used for the quantification of aligned reads. Finally, R package DEseq2 (Love et al., 2014) was used for the analysis of differential gene expression and normalization. For volcano plots, R package ggplot2 (Ito and Murphy, 2013) was used. Pathway and enrichment analysis of differentially expressed genes was carried out with the R package clusterProfiler (Wu et al., 2021).

## 3 Results

### 3.1 Plant growth-promoting traits of *P. fluorescens* and *Serratia* species

We investigated six bacterial strains with biological control activity, *P. fluorescens* SLU99, *Serratia rubidaea* EV23, AV10, *Serratia plymuthica* AS13, S412, and *Serratia proteamaculans* S4. To evaluate the growth promotion activity of these bacteria, potato plants grown in field conditions were subjected to treatment with these bacteria. Four out of the six tested strains, SLU99, EV23, AV10 and S412 resulted in significantly increased plant height, compared to the water treated control (Figure 1).

**Fig. 1.**
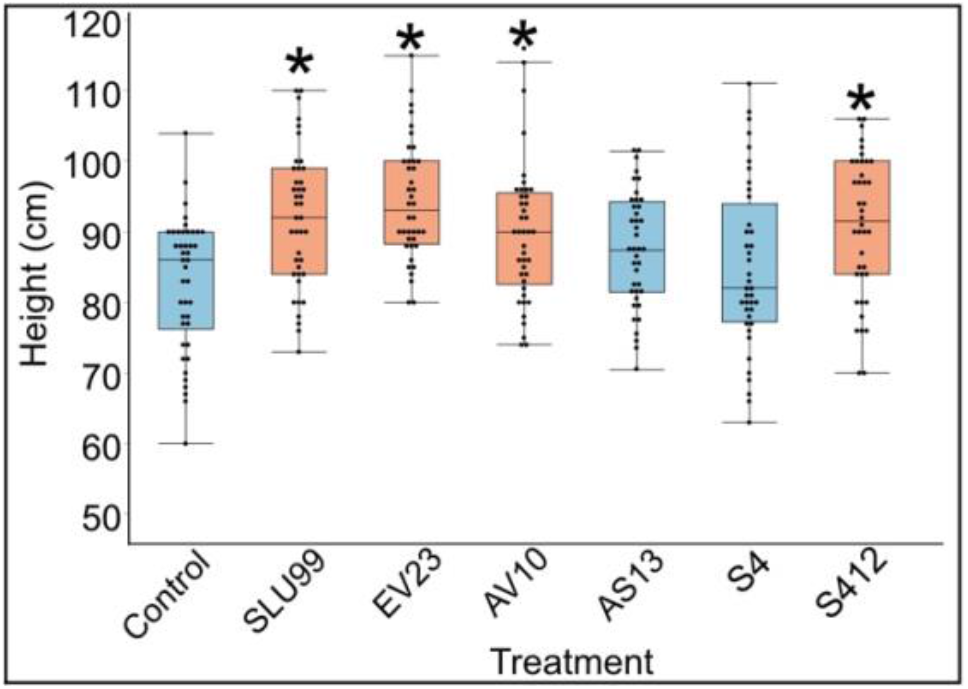
Effects of bacterial inoculations on the height of potato plants grown under field conditions. Strains used were *Pseudomonas fluorescens* SLU99, *Serratia rubidaea* EV23 and AV10, *S. plymuthica* S412 and AS13, and *S. proteamaculans* S4. Asterisks indicated statistical significance (* p < 0.001).

To examine the effectiveness of plant growth promotion, potato and tomato plants grown under greenhouse conditions were inoculated with the four selected strains. Inoculation with *P. fluorescens* SLU99 significantly increased plant height, total dry weight, and yield of tomato plants by 13.3%, 34.5%, and 183.9%, respectively, compared to non-inoculated plants. Meanwhile, *S. rubidaea* AV10-treated tomato plants had 12.3% greater total dry weight than non-inoculated plants. On the other hand, inoculation with *S. rubidaea* EV23 and *S. plymuthica* S412 had no significant effect on all morphological parameters tested on tomato plants (Table 1, Figure 2A-F). Furthermore, inoculation with both SLU99 and EV23, individually, significantly increased potato plant height, yield, and total plant dry weight (Table 1). Inoculation with AV10 promoted plant growth as total plant dry weight but not plant height, and vice versa for S412 on potato. Nevertheless, all inoculated plants had significantly higher tuber yields compared to the controls. The highest increase in yield for potato was recorded for EV23 (15%) and SLU99 (13.7%), followed by AV10 and S412 treated plants with 11.3%, and 10.7%, respectively (Table 1).

**Table 1.** Plant height at 30 and 60 days after transplanting, yield, and total dry weight of tomato and potato plants inoculated with *P. fluorescens* SLU99, *Serratia rubidaea* EV23, AV10, *S. plymuthica* S412 and control (no inoculation).

| Treatment | Plant height at 30 DAT (cm) |  | Plant height at 60 DAT (cm) |  | Yield/plant (g) |  | Total plant dry weight (g) |  |
| --- | --- | --- | --- | --- | --- | --- | --- | --- |
|  | Tomato | Potato | Tomato | Potato | Tomato | Potato | Tomato | Potato |
| *P. fluorescens* SLU99 | 44.43 | 62.78 | 70.50<sup>a</sup> | 66.80<sup>a</sup> | 73.3<sup>a</sup> | 149.0<sup>a</sup> | 44.8<sup>a</sup> | 48.5<sup>a</sup> |
| *S. rubidaea* EV23 | 42.30 | 62.44 | 59.57<sup>c</sup> | 66.12<sup>a</sup> | 32.0<sup>b</sup> | 150.7<sup>a</sup> | 34.6<sup>c</sup> | 47.8<sup>a</sup> |
| *S. rubidaea* AV10 | 47.83 | 59.00 | 65.90<sup>ab</sup> | 62.81<sup>bc</sup> | 30.7<sup>b</sup> | 145.8<sup>a</sup> | 37.3<sup>b</sup> | 47.2<sup>a</sup> |
| *S. plymuthica* S412 | 46.13 | 59.70 | 63.77<sup>bc</sup> | 64.76<sup>ab</sup> | 30.3<sup>b</sup> | 145.0<sup>a</sup> | 35.2<sup>bc</sup> | 46.4<sup>ab</sup> |
| Control | 45.57 | 58.37 | 62.20<sup>bc</sup> | 62.10<sup>c</sup> | 25.8<sup>b</sup> | 131.0<sup>b</sup> | 33.3<sup>c</sup> | 43.8<sup>b</sup> |
| *p* value | 0.2420 (ns) | 0.0709 (ns) | 0.0131 (*) | 0.0005 (**) | <0.0001 (***) | 0.0103 (*) | <0.0001 (***) | 0.0152 (*) |

**Fig. 2.**
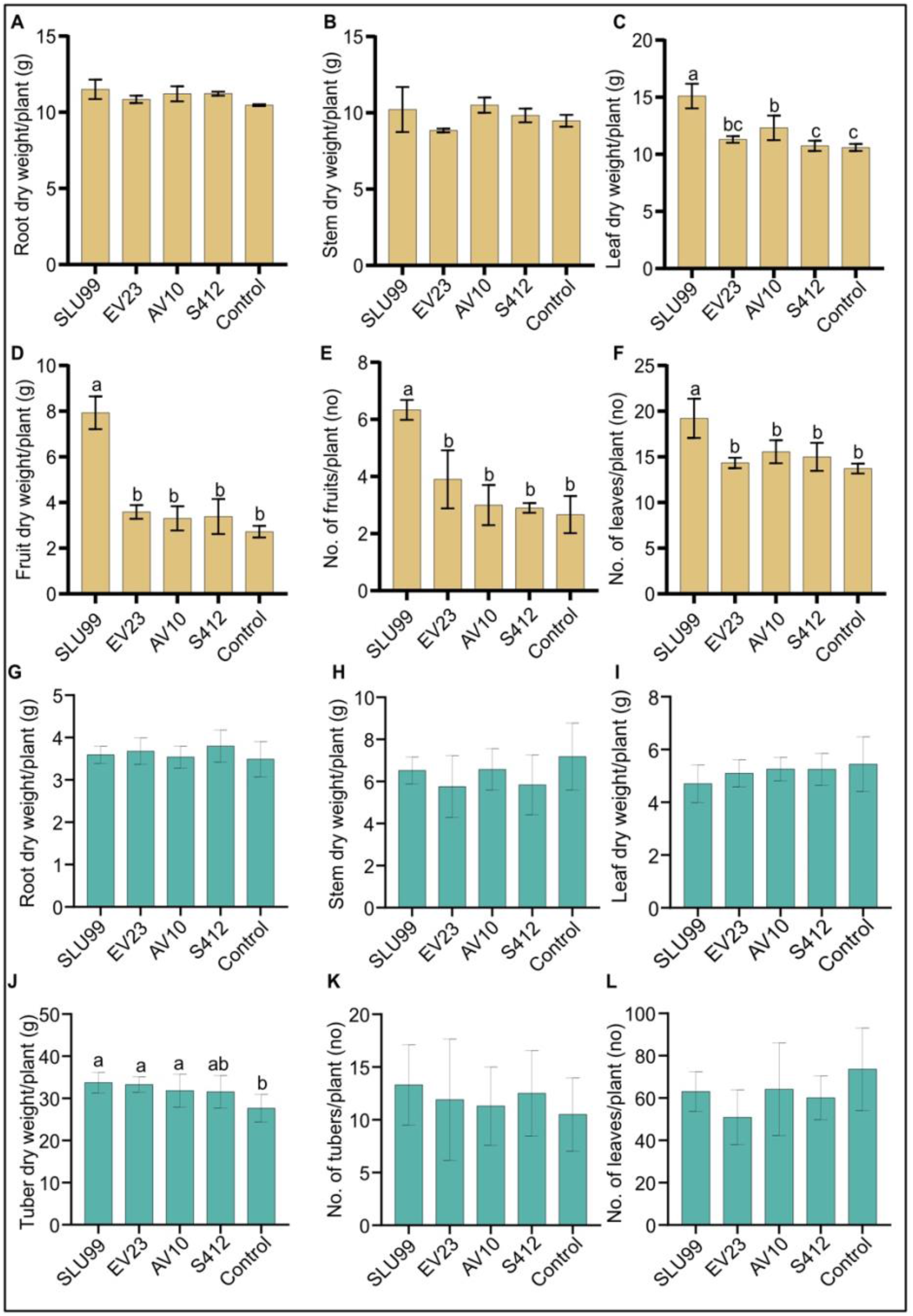
Effects of bacteria inoculations on the phenotypic characteristics of tomato and potato plants. Dry weight of different tomato (A-D) and potato (G-J) plant parts, and fruit (E), tuber (K) and leaf (F, L) number per plant. Means of root and stem dry weight were not significantly different among treatments. Strains used were *Pseudomonas fluorescens* SLU99, *Serratia rubidaea* EV23 and AV10, *S. plymuthica* S412. Means within the same bar graph without a common letter are significantly different by LSD’s test at 95% confidence level. Each value is the mean of nine and ten biological replicates per treatment for tomato and potato plants, respectively. Error bars represent standard deviations (SD).

Plant height increased rapidly during the vegetative stage until fruit set (40 DAT), or until the start of tuber initiation (30 DAP) for tomato and potato plants, respectively, and was not significantly different among treatments. Thereafter, during the fruit growth stage (50-60 DAT), a significant effect of bacteria inoculation on tomato plant height was observed (P=0.0431 and P=0.0131 at 50 and 60 DAT, respectively) (Table 1, Supp Figure 1B). In contrast, a 4.2% reduction in tomato plant height was observed when plants were treated with EV23 (Table 1). Applications of SLU99 resulted in increased tomato leaf (Figure 2E) and fruit number (Figure 2F) compared to other treatments. Consequently, SLU99-treated tomato plants had 39.7% and 191.2% more leaf and fruit dry weights, respectively, compared to non-inoculated plants.

For potato plants at 30 DAP, swelling stolon tips were observed in all treatments suggesting tuber initiation had started. The effect of bacteria inoculation on potato plant height was only significant during the tuber bulking stage (40-60 DAP) (Table 1, Supp Figure 1). In addition, all bacteria except S412 significantly increased tuber dry weight over control plants (P=0.0003). However, the dry weight of other plant parts was not enhanced by bacteria treatments (Figure 2G-I).

Relative chlorophyll content was significantly higher after SLU99 application in tomato (P=0.0188) and potato plants (P=0.0287) but was not improved after EV23, AV10 and S412 treatments compared to the control (Figure 3).

**Fig. 3.**
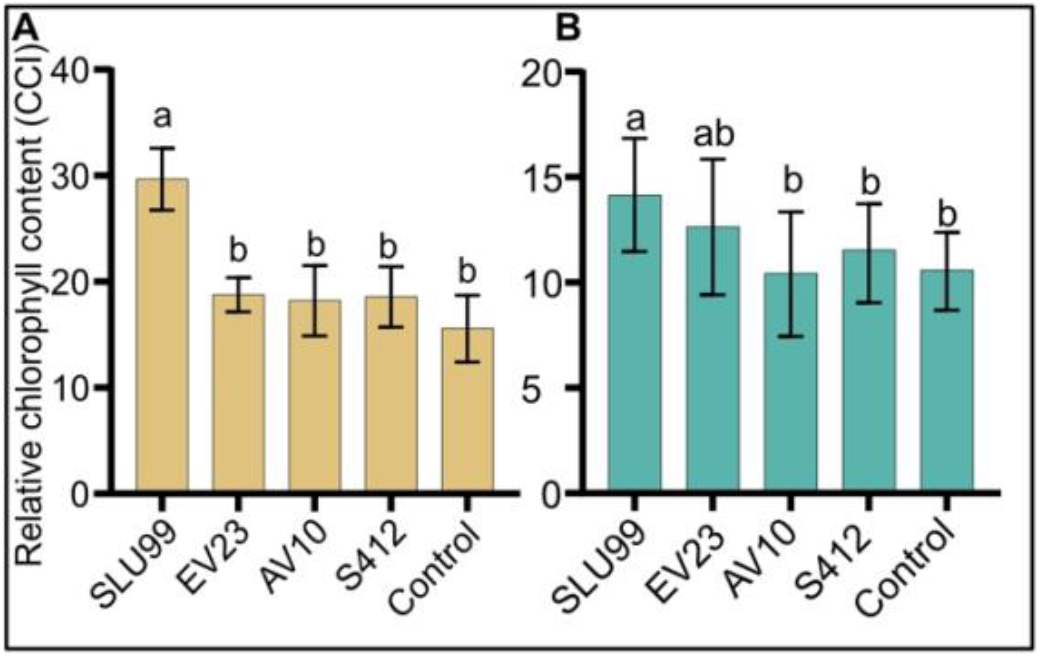
Relative chlorophyll content of tomato and potato plants inoculated with rhizobacteria. The values presented are mean values of relative chlorophyll content in the unit of Chlorophyll Content Index (CCI) measured using an Apogee MC-100 Chlorophyll Concentration Meter at harvest. Each value is the mean of 90 readings from nine biological replicates for tomato and 100 readings from ten biological replicates for potato per treatment. Means within the same bar graph without a common letter are significantly different by LSD’s test at 95% confidence level.

Additionally, at harvest, inoculation with SLU99 resulted in 3.4% higher total nitrogen (TN) in the tomato grown soil, while inoculation of EV23, AV10 and S412 resulted in 10.7%, 7.7% and 24.3% lower TN over control treatment, respectively (Supp. Table 1). The elements of P & K were slightly increased in tomato grown soil after SLU99 treatment, whereas EC and SOM were decreased. In potato grown soil, all bacterial treatments increased available K over the control by 22.2%-66.7%. TN in potato grown soil decreased in a small range after SLU99, EV23 and AV10 treatments (2.1%-6.2%). Meanwhile, the available P, pH, EC, and SOM at harvest of soils grown with potato were not affected by bacteria treatments (Supp. Table 1). In summary, inoculation of SLU99 significantly promoted the growth of tomato and potato plants, and EV23 promoted the growth of potato plants grown in a controlled environment as reflected by high chlorophyll content, total plant biomass and yield.

### 3.2 Effects of rhizobacteria inoculation on growth of *in vitro* grown tomato and potato plants

To gain insights into the mechanisms behind growth promotion, tomato and potato plants were grown in MS media supplemented with the sterile-filtered bacterial culture supernatant. When compared to the mock treatment, tomato seeds grown in the SLU99-supernatant supplemented media showed a higher germination rate, increased root growth and shoot height, and enhanced number of lateral roots (Figure 4A, B). Potato explants grown in the media supplemented with SLU99 and EV23 supernatant resulted in increased shoot height by 24.6 % (*p* = 0.02) compared to the control (Figure 4C, D). SLU99 treatment also increased root growth and resulted in the formation of adventitious roots and secondary adventitious roots (SAR), which emerge from aerial parts (Figure 4D).

**Figure 4.**
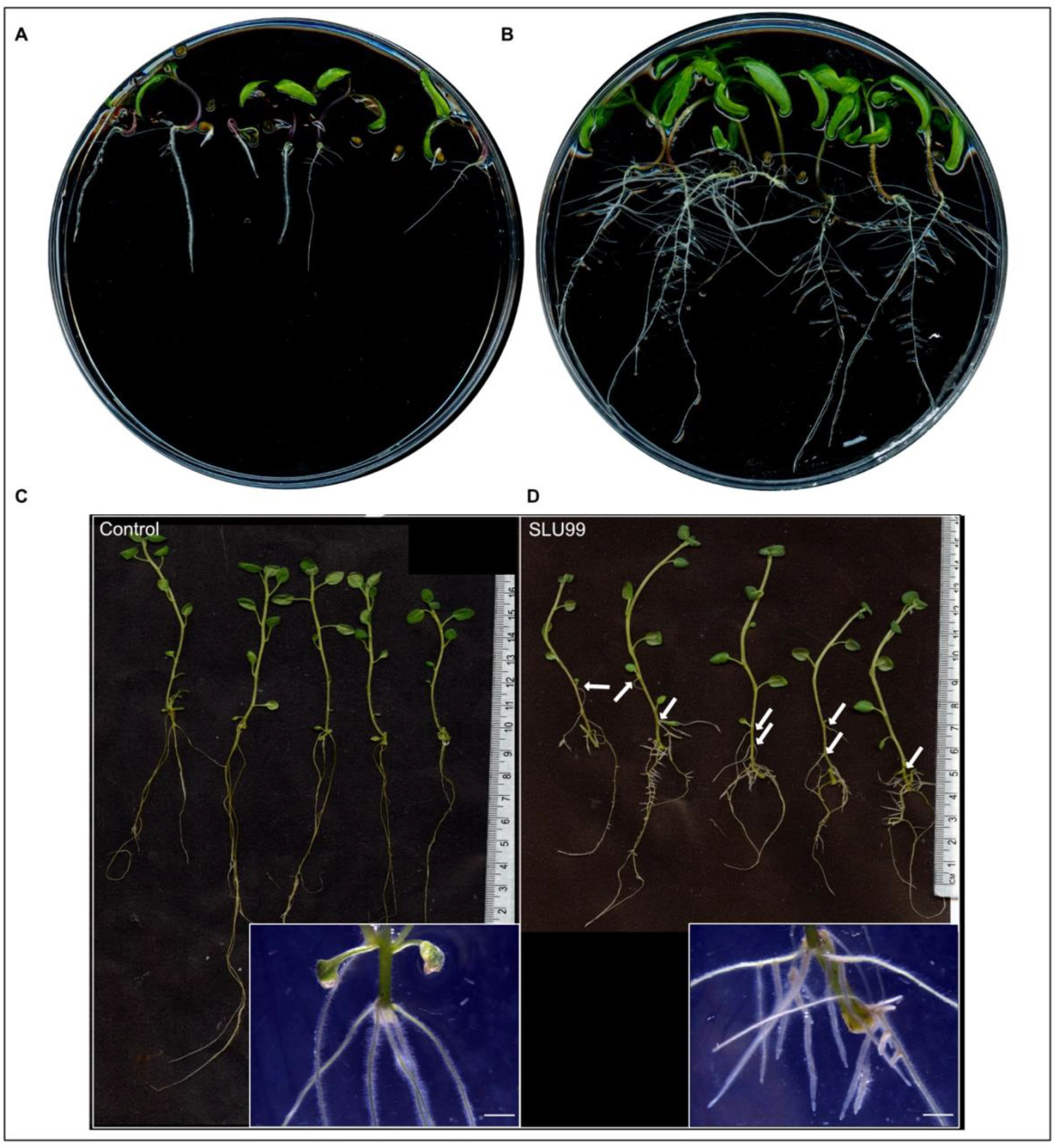
Effect of *Pseudomonas fluorescens* SLU99 on plants grown *in vitro*. Growth of tomato seedlings upon treatment with culture supernatant of SLU99 (B) compared to mock treatment (A). Growth of potato plants treated with *P. fluorescens* SLU99 (D) compared to control treatment (C). Arrows indicate secondary adventitious roots developed upon treatment with SLU99. Insets in C and D are representative stereo microscope images at root induction. The scale bar in the inset images represents 1 mm.

### 3.3 Differentially expressed genes (DEGs) and KEGG pathway enrichment analysis

To gain insights into the mechanisms of SLU99-mediated growth promotion in tomato and potato plants, RNA extracted from the roots and leaves grown in the media supplemented with the supernatant of SLU99 was subjected to transcriptome profiling. A total of 1193 and 2226 genes were differentially expressed (log2FC >1.5, *p* < 0.05) in the roots of tomato and potato, respectively, compared to the control samples (Table 2). Among them, 1076 and 996 genes were upregulated, and 117 and 1230 genes were downregulated in the respective samples (Table 2, Fig. 5A and B). In tomato leaves, a total of 1732 DEGs were detected (log2FC >1.5, *p* < 0.05) with 1322 genes upregulated and 410 genes downregulated (Fig 5C). For potato leaves, we identified 959 (log2FC >1.5, *p* < 0.05) when treated with SLU99 compared to the control samples. Of the identified DEGs, 206 were upregulated, and 753 genes were downregulated (Table 2 and Fig 5D).

**Table 2.** Characteristics of the DEGs in tomato and potato plants treated with *Pseudomonas fluorescens* SLU99 culture supernatant.

|  | Tomato |  | Potato |  |
| --- | --- | --- | --- | --- |
|  | Root | Leaf | Root | Leaf |
| **Upregulated** | 1675 | 2024 | 1787 | 411 |
| Log2FC > 1.5 | 1094 | 1467 | 1012 | 209 |
| p value < 0.05 | 1076 | 1322 | 996 | 206 |
| **Downregulated** | 319 | 841 | 2253 | 1112 |
| Log2FC < -1.5 | 118 | 420 | 1251 | 772 |
| p value < 0.05 | 117 | 410 | 1230 | 753 |

**Figure 5.**
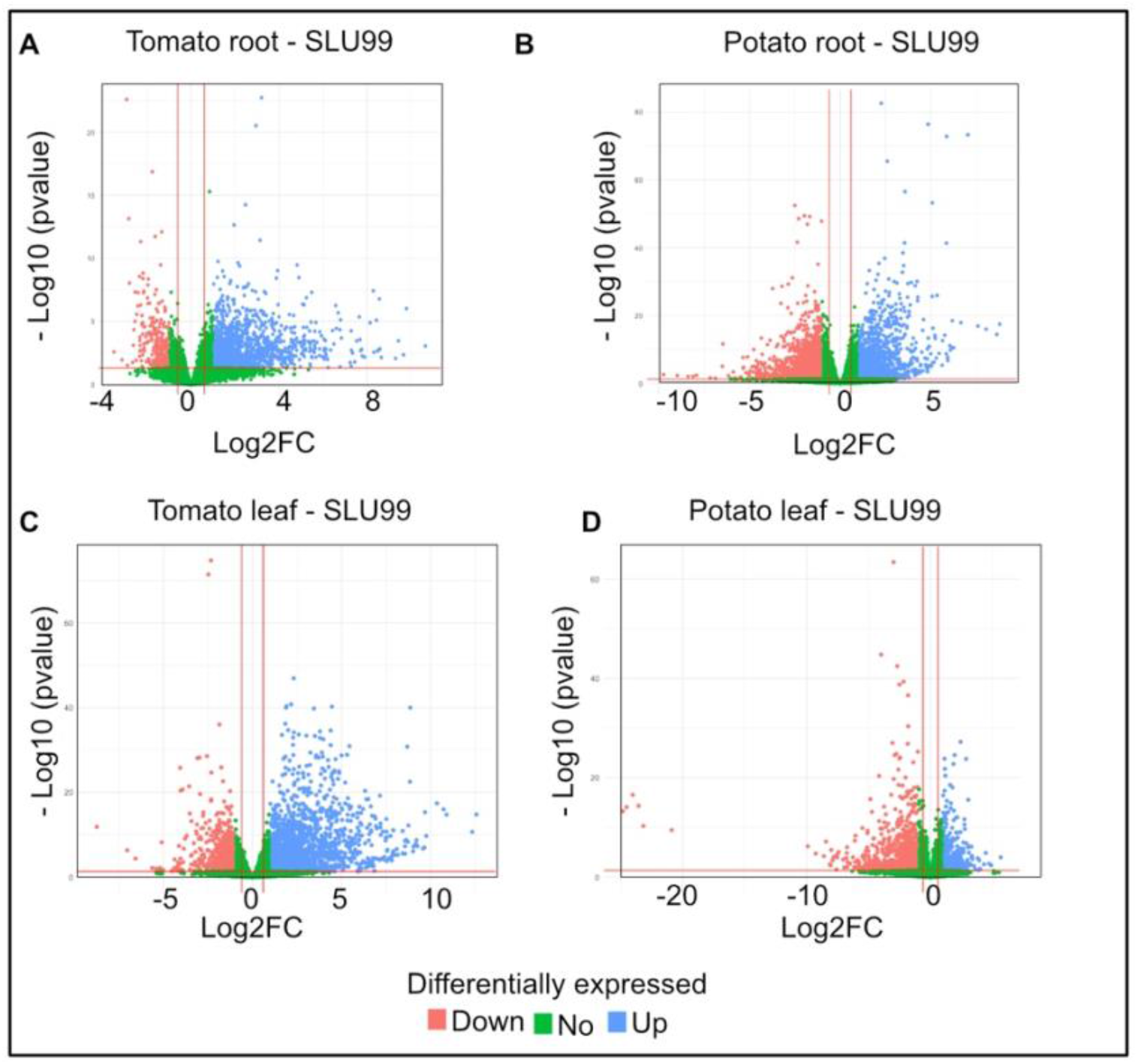
Volcano plot of differentially expressed genes (DEGs) for potato and tomato leaves and roots treated with the culture supernatant of *Pseudomonas fluorescens* SLU99, compared to the mock treatment. Upregulated genes are shown as blue dots at the right side of each plot; downregulated genes are shown as pink dots at the left side of each plot; non-differentially expressed genes are shown as green dots clustered at the centre (centred around Log2FC 0) of each plot.

We performed KEGG pathway enrichment analysis to gain a better understanding of the functional categories of the DEGs in response to the PGPR culture supernatant treatment. In the treated root samples, a total of 300 and 753 genes were grouped into enriched pathways in tomato and potato, respectively, and a total of 382 and 259 genes were grouped into enriched pathways in the treated leaves of tomato and potato, respectively. Treatment resulted in enrichment for genes involved in plant hormone signal transduction pathway (37 and 33 genes, respectively), zeatin biosynthesis (15 and 13 genes), plant-pathogen interactions (37 and 25 genes) and MAPK signalling pathway (30 and 19 genes) in both tomato and potato leaves. Additionally, a total of 40, 76, 44 and 25 genes related to the phenylpropanoid biosynthesis pathway were enriched in the roots and leaves of tomato and potato upon treatment with SLU99 culture supernatant (Fig 6).

**Fig. 6.**
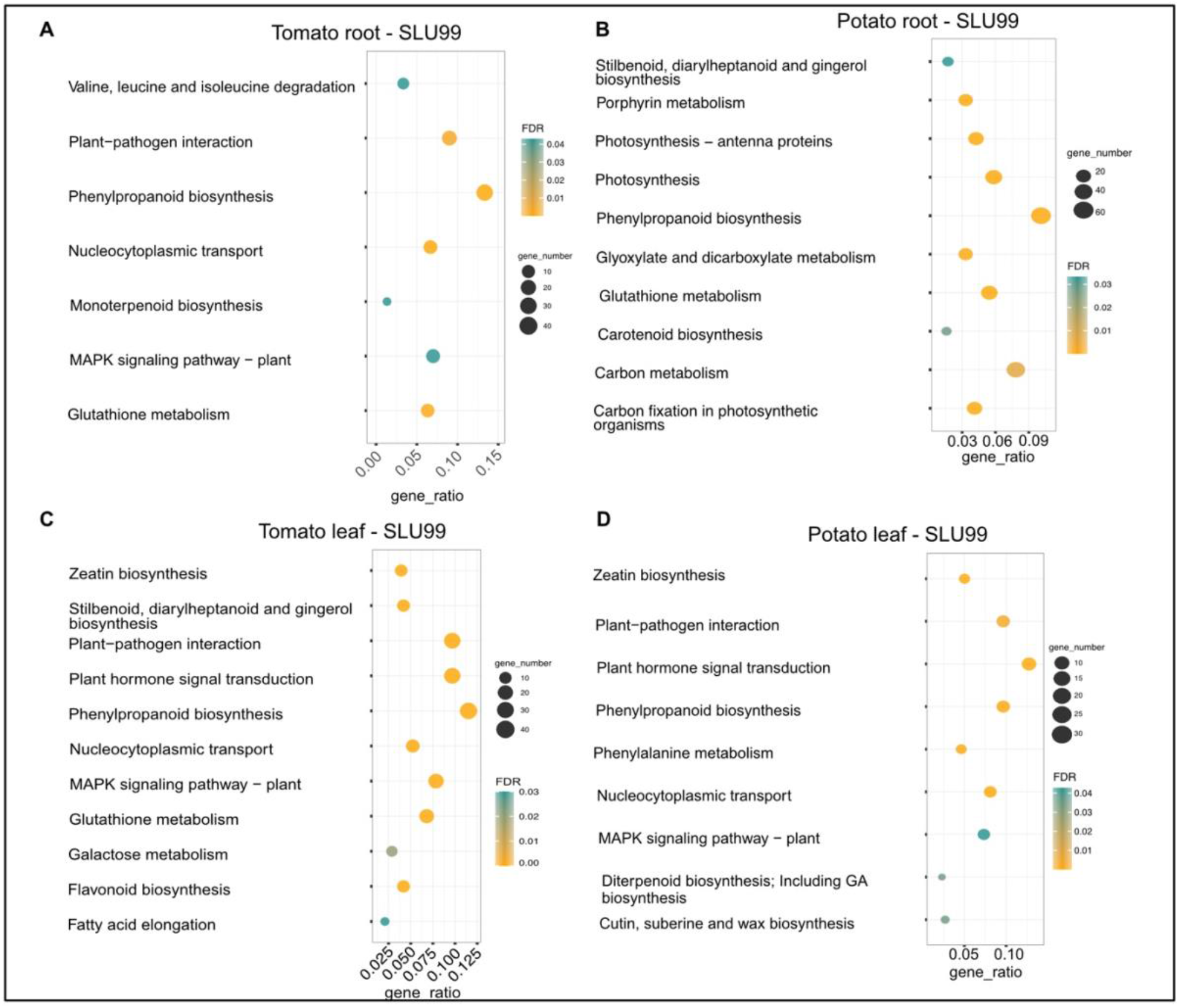
KEGG enrichment analysis of DEGs in the transcriptomes of potato and tomato leaves and roots treated with the culture supernatant of *Pseudomonas fluorescens* SLU99, compared to the mock treatment. The size of the circles in each plot represents the number of DEGs annotated for that pathway or process. The gene ratio on the x-axis represents the ratio of the count of core enriched genes to the count of pathway genes.

To compare the effects of the tomato and potato samples, we examined the overlap of the DEG products among the roots and leaves. The analysis showed an overlap of 164 gene products in the root samples, and 95 in the leaf samples upon SLU99 treatment in potato and tomato plants (Fig. 7A and B). Within the species, there was an overlap of 515 gene products between root and leaf samples of tomato, while 265 similar gene products were responsive in the root and leaf samples of potato (Fig. 7C and D).

**Fig. 7.**
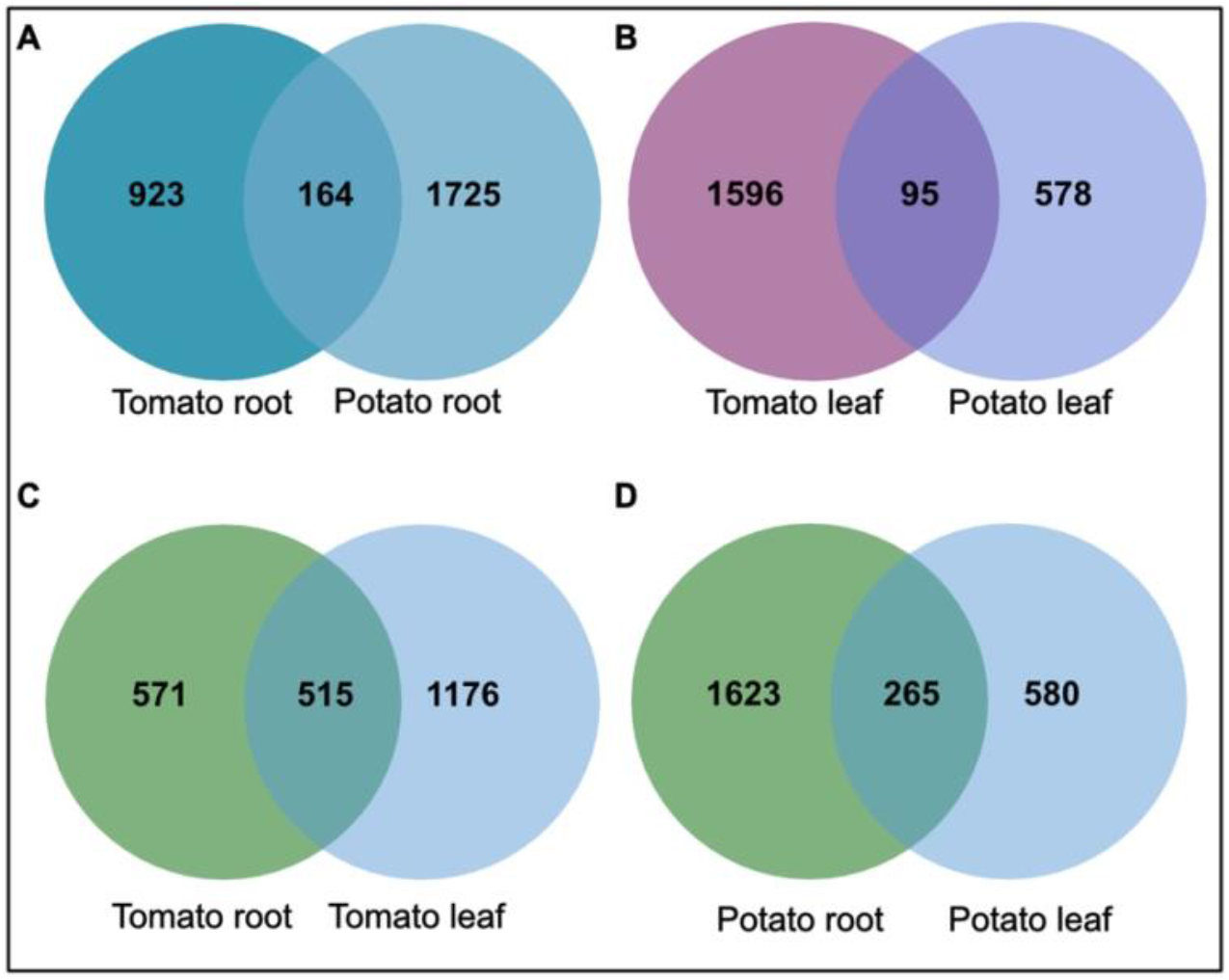
The overlap of products of differentially expressed genes (DEGs) in potato and tomato leaves after treatments with *Pseudomonas fluorescens* SLU99 supernatant.

### 3.4 SLU99 regulates phytohormonal biosynthesis and signal transduction pathways

One of the major objectives of this study was to elucidate the changes in the host transcriptomes that are important for PGP activity. Phytohormones, as a result of their complex interaction and crosstalk, regulate various cellular processes involved in plant growth and development. The gibberellins (GAs) play an important role in several developmental processes including seed germination. Upon SLU99 treatment in tomato root, *gibberellin 3-beta-dioxygenase 1*-like (*GA3OX1*), a gene involved in the biosynthesis of GA was upregulated (log2FC 1.9, *p*<0.05). On the other hand, in tomato leaves, *gibberellin 2-beta-dioxygenase 8* (*GA2ox8*), a gene involved in the deactivation of GA, was upregulated (log2FC 1.6, *p*<0.01) (Fig.8).

**Figure 8.**
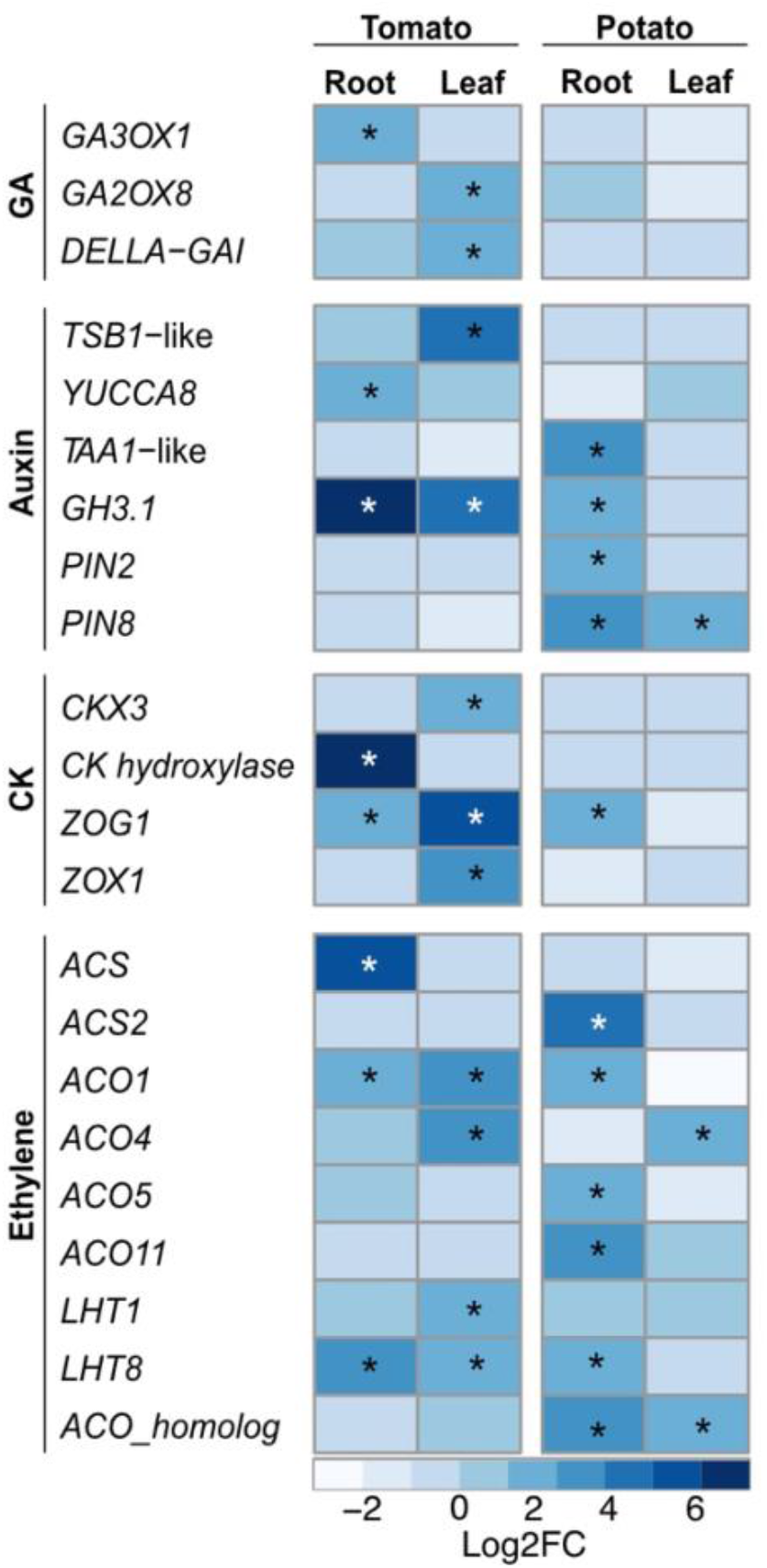
Heatmap of key differentially expressed genes (DEGs) involved in different phytohormonal pathways upon treatment with *Pseudomonas fluorescens* SLU99 culture supernatant. GA = gibberellic acid; CK = cytokinin. Asterisks indicated statistical significance (* p < 0.05).

The hormones auxin and CK are major players in the regulation of signaling pathways underlying plant growth and development. In tomato, *tryptophan synthase beta subunit 1* (*TSB1*)-like, responsible for increased in tryptophan biosynthesis, and *YUCCA8*, responsible for the last enzymatic step from indole-3-pyruvic acid (IPA) to IAA were upregulated by 16 and 3 times (log2FC 4 and 1.6) in leaf and root samples, respectively. Additionally, in potato roots, SLU99 treatment resulted in increased expression (log2FC 3.3) of *L-tryptophan-pyruvate aminotransferase 1 (TAA1)-*like, which encodes an enzyme that converts L-tryptophan to IPA (Fig. 8).

The activation of auxin biosynthesis genes and production of IAA by SLU99 (Kelbessa et al., 2022) prompted us to analyze the differences in the expression of genes involved in auxin signaling pathways. Major auxin-responsive genes such as *Gretchen Hagen 3* (*GH3*) and *small auxin up RNA* (*SAUR*) are transcriptionally regulated by auxin at early stages of signal transduction. Following treatment with SLU99 culture supernatant, expression of the *IAA-amido synthetase GH3*.*1 (GH3*.*1)*, important for maintaining auxin homeostasis increased significantly (log2FC 6.8, 4.6 and 1.6 in tomato root, tomato leaf and potato root, respectively) in comparison to the mock treatment. Within the SAUR family, five and seven genes were differentially expressed in the tomato and potato plants, respectively. Additionally, *PIN2*, a root-specific auxin transporter and *PIN8*, a constitutively active auxin transporter were also upregulated in the potato plants (Fig.8).

The gene encoding CK hydroxylase, an enzyme that catalyzes the biosynthesis of trans-zeatin (tZ), a biologically active CK, was strongly upregulated (log2FC 7.5) upon treatment with SLU99 in tomato roots. On the other hand, in tomato leaf *CK dehydroxygenase 3* (*CKX3*), involved in the degradation of CK was upregulated 4-fold (log2FC 2) compared to the mock treatment. Furthermore, *zeatin O-glycosyltransferase* (*ZOG1*) and *zeatin O-xylosyltransferase* (*ZOX1*) were upregulated in tomato leaves (log2FC 5.4 and 3.5), while only *ZOG1* was upregulated in the tomato and potato roots (log2FC 2.1 and 1.6). *ZOG1* and *ZOX1* are involved in the conversion of tZ to a stable and reversible O-glycosylzeatin and O-xylosylzeatin, respectively.

The PGP activity of bacteria is also attributed to the changes in the plant hormone ethylene (Poupin et al., 2016). Since SLU99 displayed ACC deaminase activity, we examined changes in the expression of genes encoding the enzymes of the ethylene biosynthesis pathway. *1-aminocyclopropane-1-carboxylate_synthase-*like (*ACS*) involved in the synthesis of ACC, a direct precursor of ethylene, and *1-aminocyclopropane-1-carboxylate_oxidase* (*ACO*) subsequently oxidizes ACC to ethylene. In tomato, *ACS* and *ACO1* were upregulated in roots (Log2FC 5.1 and 1.8, respectively), while *ACO1* and *ACO4* were upregulated in leaves (Log2FC 3.2 and 2.6, respectively). In potato, *ACS2* was upregulated in roots (log2FC 4.1), while *ACS4* was upregulated in leaves (log2FC 2.2). *ACO* homologs were upregulated both in the roots and leaves (log2FC 2.9 and 1.7, respectively), while *ACO1, ACO5* and *ACO11* were upregulated in leaves (log2FC 2.1, 2.1 and 3.3, respectively). Lysine histidine transporters (LHT) are associated with the transportation of ACC. When treated with SLU99 culture supernatant, *LHT8* was upregulated in both tomato and potato roots and tomato leaves by 3.6, 2.5 and 2.5 times (log2FC), respectively, while *LHT1* was upregulated only in tomato leaves (log2FC 1.6). Taken together, these results indicate that treatment with the culture supernatant from SLU99 stimulates the accumulation of phytohormones auxin, CK and ethylene.

### 3.5 Reprogramming of host transcriptional networks

The discovery of several DEGs involved in signal transduction pathways in response to SLU99 treatment suggests strong regulation of host transcription factor networks. To investigate the interaction of SLU99 with the plant transcriptional network, DEGs encoding TFs were identified and assigned to families using the plant transcription factor database. In total, DEGs belonging to 24 different TF families were identified with the greatest number of DEGs identified in the root samples compared to the leaf samples (Supp. Table 2). In the tomato and potato roots, a total of 112 and 182 TF encoding DEGs were found, whereas 62 to 92 DEGs were found in leaf samples, implying significant reprogramming of the host transcriptional network. The HD-ZIP TF genes were exclusively upregulated in tomato plants (11 upregulated and 2 downregulated), while 14 HD-ZIPs were downregulated in potato. Notably, the strongest upregulation was found for *ATHB12* (log2FC 9.6) followed by *ATHB40-like* (log2FC 6.4), and *ATHB7-like* (log2FC 3.4) in tomato leaves. *ATHB12* and *ATHB7*, considered paralogs, belong to the class-I HD-ZIP TF family and were also upregulated in root samples (log2FC 2.1 and 2.0, respectively). *ATHB40*, on the other hand, is a class-II HD-ZIP TF that was upregulated in the root (log2FC 3.7). *ATHB12* and *ATHB7* are associated with root elongation and leaf development, albeit at different stages (Ré et al., 2014; Hur et al., 2015). *ATHB40* is a negative regulator of primary root development (Mora et al., 2022).

In addition to the PGP-related TF encoding DEGs, we also identified several DEGs encoding for TFs involved in defence-related pathways, namely, WRKY, MYB, MYC, HSF, and NAC TFs. WRKY TFs are global regulators of host responses to phytopathogens. Treatment with SLU99 culture supernatant triggered differential expression of 14 WRKY genes in tomato and 11 genes in potato. Two of the WRKY TF encoding genes, *WRKY30* and *WRKY45*, involved in defence against biotic and abiotic stresses, were upregulated in common between tomato and potato. *WRKY6, WRKY55*, and *WRKY71* were uniquely upregulated in tomato leaf and, along with *WRKY33B* and *WRKY75*, are involved in defence against necrotrophic fungal pathogens. *WRKY40* and *WRKY41*, genes involved in abiotic stress tolerance, were strongly upregulated in tomato leaves (log2FC 9.0 and 4.5, respectively), and were downregulated in potato leaves upon SLU99 treatment (log2FC -2.1 to -2.7). The *MYC2* TF upregulated (log2FC 5.1) in tomato roots upon SLU99 treatment indicates a role in JA-mediated induced systemic resistance (ISR). Furthermore, treatment with SLU99 induced expression of several R-gene orthologs such as *Cf-2,2-like* (log2FC 2.2), *Cf-9* (log2FC 3.4), *RPM1* (log2FC 1.7), *RPS5* (log2FC 4.1), *CSA1* (log2FC 4.8) and *Alternaria stem canker resistance* (log2FC 1.6) in tomato leaves, *LEAF RUST 10 DISEASE-RESISTANCE LOCUS RECEPTOR-LIKE PROTEIN KINASE-like 1*.*1* (*LRK10*) (log2FC 2.9 and 5.3 in root and leaf, respectively), *TMV resistance protein N* gene (log2FC 6.0) in tomato root and *putative late blight resistance protein homolog R1A-10* (log2FC 1.6) and *R1A-3* (log2FC 1.5) in potato leaves.

## 4 Discussion

Several soilborne rhizobacteria have long been known to promote plant growth across a wide range of plant species. In this study, six bacterial strains were chosen for their biocontrol potential and tested on potato plants in a late-blight hotspot field. Four of the strains that directly impacted the height of the potato plant in the field were chosen for detailed examination in a controlled environment to further evaluate their growth-promoting function. These strains included *P. fluorescens* strain SLU99, *S. rubidaea* strains EV23, AV10, and *S. plymuthica* S412.

In the growth chamber, all four strains enhanced at least two growth variables in potato plants, indicating that all tested strains significantly impact potato growth, which is consistent with the field evaluation. However, the effect of these strains on the dry matter accumulation is only noticeable in the tuber, which accounts for more than 60% of the total dry weight, suggesting that PGPR treatment promotes photosynthate translocation into tubers. In contrast, growth-promotion activity in tomato plants is strain-dependent with SLU99 and AV10 improving plant height and total dry weight. Furthermore, SLU99 improved chlorophyll content, total leaf number and leaf area, suggesting enhanced light interception and photosynthesis rate explaining the obtained higher tomato fruit yield.

Intriguingly, although all four strains isolated from tomato rhizospheres are favorable to potato yield, only SLU99 is significantly beneficial to tomato yield. Previous research found that bacterial consortiums aid plant growth by enhancing stress tolerance (Silambarasan et al., 2019; Yang et al., 2021; Kelbessa et al., 2023). Recently, results from our group demonstrated that SLU99 is compatible with strains EV23 and AV10 (Kelbessa et al., 2022), implying that these bacteria could co-exist as a consortium in the rhizosphere. Considering that all these strains are potent biocontrol agents (Kelbessa et al., 2022), a bacterial consortium of these strains could act synergistically to contribute to reducing disease under natural conditions. Moreover, plant exudates attract rhizobacteria to colonize the roots (Bais et al., 2006). It has been demonstrated that the composition of root exudates varies between cultivars of the same species (Zhang et al., 2020) and thus cultivar-specific traits might have contributed to a negative regulatory effect of the colonization of *Serratia* species in the tomato variety used in this study.

PGPR also enhances the mobilization of locally available nutrients for plant uptake by solubilizing P, K, and Zn, and nitrogen fixation (Prasanna et al., 2016). N and K are the most abundant nutrients in plant tissues (Sardans and Peñuelas, 2015). Potato cultivation takes up K in large quantities and is very important to gain a higher yield of marketable tubers (Khan et al., 2012). Our results suggest increased K in the soil following the treatment with PGPR, consistent with a previous study that showed *Enterobacter cloacae*, a PGPR, led to a higher amount of K in the soil (Ghadam Khani et al., 2019). However, further research validation is required to assess the impact of PGPR on soil nutrient contents. On the other hand, in tomato, TN, available P and K were slightly increased after SLU99 treatments, indicating that this strain may have facilitated nutrient availability for plant uptake (Meena et al., 2014).

Phytohormone-mediated signal transduction and their interplay regulate several physiological processes in plant growth and development. Phytohormones also mediate cellular responses during abiotic and biotic stress. Rhizobacterial-stimulated plant growth is intrinsically linked to the production of phytohormones, siderophores, and secretion of ACC-deaminase that reduces ethylene biosynthesis (Yang et al., 2009). Accordingly, KEGG pathway enrichment analysis suggested that treatment with SLU99 culture supernatant resulted in significant differential expression of genes involved in plant hormone signal transduction. Genes involved in the biosynthesis of zeatin, a naturally occurring CK that promotes cell division in plants, were significantly differentially expressed in both potato and tomato leaves upon SLU99 treatment. Strain SLU99 enhanced the expression of *CK hydroxylase* in tomato root suggesting an increase in the biosynthesis of trans-zeatin, which is reported to be transported through the xylem (Osugi et al., 2017). However, excess CK needs to be stored to be protected against CK oxidases. Conversion of zeatin to O-glucosyl- and O-xylosyl-zeatin, a reversible process, is important for the storage of CK. Our results show that *ZOG1* and *ZOX1*, genes that encode enzymes in the glycosylation of zeatin are upregulated, suggesting the CK is synthesized in excess and is stored in roots and leaves, upon SLU99 treatment in tomato.

The increase in the germination of the tomato seeds requires specific reprogramming of the GA pathway (Groot and Karssen, 1987). Moreover, the presence of GA in the root meristem and elongation zone is necessary for the normal growth of the root. *GA3OX* encodes a key enzyme involved in the last step in the biosynthesis of the GA and is reported to be expressed in the root elongation zone (Barker et al., 2021). SLU99 culture supernatant treatment of tomato plants resulted in an increased expression level of *GA3OX*, suggesting an increase in the GA in the roots. GA homeostasis is necessary for plant growth and development (Richards et al., 2003). GA 2-oxidation mediated by the *GA2ox* gene family is reported to be a major GA inactivation pathway that functions throughout *Arabidopsis* development (Schomburg et al., 2003; Rieu et al., 2008) and is necessary for maintaining GA levels in peach (Cheng et al., 2021). Our results show that *GA2ox8* is upregulated in tomato leaves following treatment with SLU99 culture supernatant, suggesting PGPR aid in maintaining GA levels. Our results of increased expression of genes encoding DELLA-GAI and a slight increase of AP2/ERF (log2FC 1.2, *p* 0.04) in tomato leaves are in line with reports showing lowered GA levels are also associated with improved stress tolerance through DELLA-mediated activation of AP2/ERF (Colebrook et al., 2014; Castro-Camba et al., 2022).

It is well-established that several auxin producing PGPR regulate auxin localization and distribution in the plant (Tsukanova et al., 2017). Rhizobacterial-produced auxin has also been shown to be important for PGPR-mediated morphological changes in plants (Spaepen et al., 2014). Treatment with these PGPR also resulted in increased levels of endogenous auxin and is associated with root growth. Indeed, local auxin maxima in the pericycle is necessary for the formation of lateral root primordia and their subsequent development into lateral roots.

However, the effect of exogenous auxin on the root is concentration dependent. At higher concentrations, exogenous auxin inhibits root growth (Ivanchenko et al., 2010). Consequently, large amounts of auxin generated by some rhizobacteria strains, such as *Enterobacter* (Park et al., 2015), *P. fluorescens* (Zamioudis et al., 2013) and *P. syringae* (Loper and Schroth, 1986) inhibit primary root elongation in lettuce, *Arabidopsis*, and sugar beet, respectively. In short, while auxin is an important contributing factor to root growth, regulating endogenous auxin levels is also crucial. *GH3* genes, upon induction by IAA accumulation, encode enzymes involved in the conjugation of free IAA to amino acids thereby maintaining auxin homeostasis (Staswick et al., 2005). Our study is consistent with earlier reports of *P. fluorescens* producing auxin and with increased lateral root formation (Chu et al., 2020; Ortiz-Castro et al., 2020). Additionally, treatment with SLU99 also impaired primary root elongation in potato plants. Taken together, our results support the hypothesis that treatment with SLU99, while increasing the endogenous IAA levels through upregulation of *YUCCA8*, and utilizing necessary auxin for lateral root development, also aided in maintaining its homeostasis through GH3.1 mediated conjugation of free auxin (Fig. 9).

**Figure 9.**
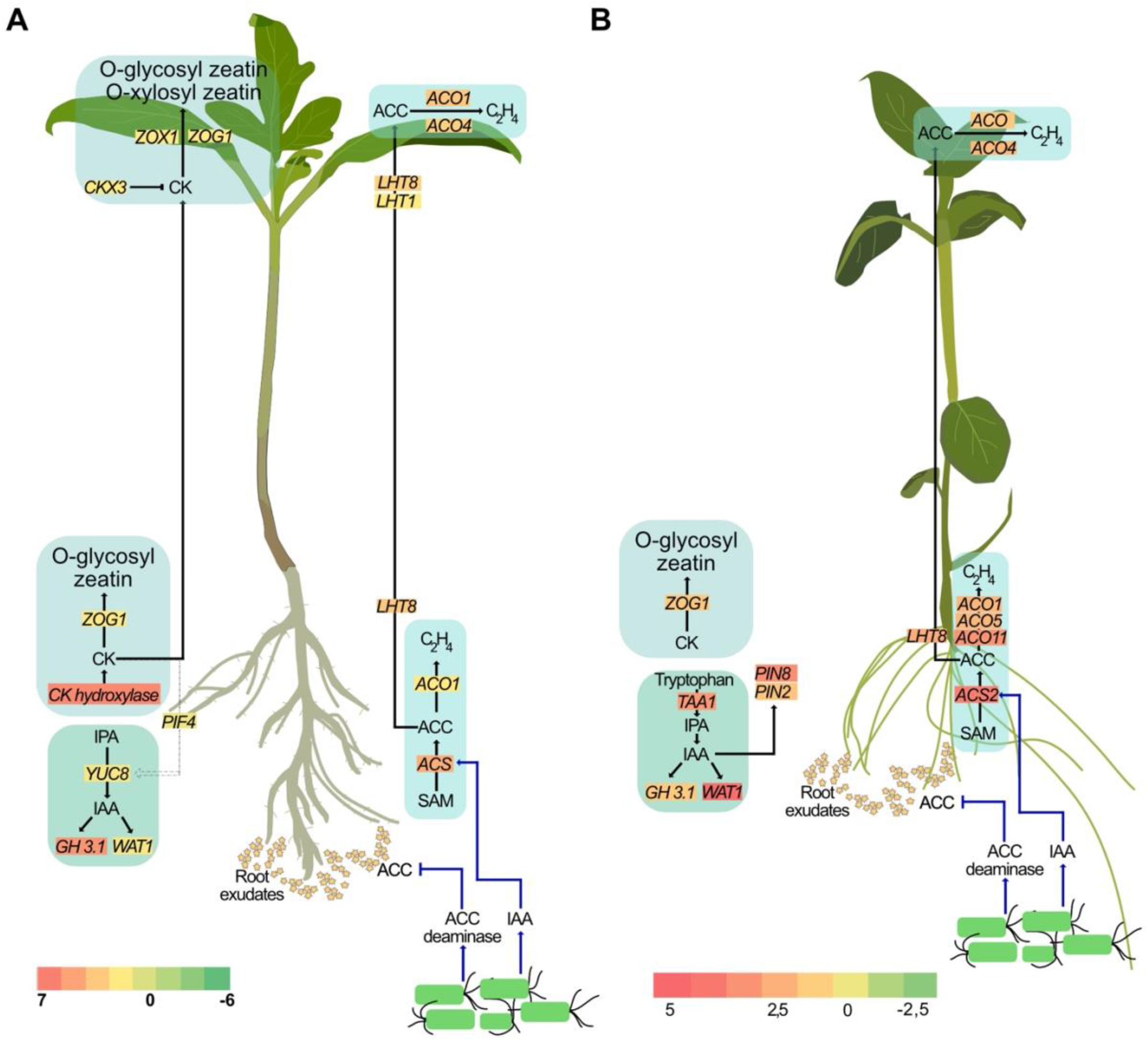
A proposed model illustrating the effect of *P. fluorescens* strain SLU99 on different phytohormonal pathways in tomato (A) and potato (B) plants.

Higher auxin levels are also associated with increased ethylene biosynthesis (Fa Yang and Hoffman, 1984). Regulating cellular ethylene content is a critical aspect of plant growth and development. Application of exogenous auxin is also reported to increase expression of *ACS* genes that encode enzymes involved in the conversion of *S*-adenosylmethionine to ACC, the ethylene precursor (Tsuchisaka and Theologis, 2004; Niu et al., 2022). Considering that PGPR produce auxin, it can be hypothesized that the auxin produced by PGPR result in increased expression of *ACS*, and thus might promote ACC production. Providing further evidence of ethylene synthesis, *ACO1*, which encodes ACC oxidase, the enzyme required for the synthesis of ethylene from ACC, is also upregulated upon SLU99 treatment. However, higher ethylene levels not only inhibit root development but also trigger an adaptive response such as growth inhibition and delayed flowering (Ravanbakhsh et al., 2018). PGPR belonging to diverse genera are reported to produce ACC deaminase, an enzyme that lowers host ACC levels (Glick, 2005). It is important to note that although bacteria produce ACC deaminase, due to the higher substrate affinity of ACC oxidase than ACC deaminase, the ethylene level in the plant cannot be totally eliminated (Glick et al., 1998). Intriguingly, two differentially expressed genes, *LHT8* and *LHT1*, belonging to the lysine histidine transporter family that was previously linked to ACC transport (Shin et al., 2015; Vanderstraeten and van Der Straeten, 2017) are upregulated in treated roots and leaves, respectively, suggesting that the excess ACC may be transported to the shoot (Fig.9). Further research using ethylene biosynthetic and signalling mutants is necessary to enhance our understanding of the involvement of PGPR in modulating ethylene levels during plant development.

Plant hormone homeostasis is critical in plant growth and development. Since PGPR can have a significant influence on hormone biosynthesis and signalling, plants need to cope with such differences. Upregulation of *CKX3, GA2OX*, and *GH3* in PGPR-treated plants in our study might thus be an adaptation mechanism to increased hormone levels.

All of the rhizobacteria strains tested here have previously been shown to have biocontrol activity against the plant pathogen *Phytophthora colocasiae* (Kelbessa et al., 2022), both in culture and *in planta*. The activation of plant metabolite pathways (growth and defence) by the rhizobacteria, and their direct antagonistic activity against pathogens, suggests that these strains will have utility in sustainable agriculture by controlling serious crop diseases and boosting yield.

## Data availability statement

All the raw high quality (Q30), adapter trimmed fastq files were deposited into the NCBI SRA database under Bioproject PRJNA851815. Accession numbers of BioSamples are SAMN29251990, SAMN29251991, SAMN29251992, SAMN29251993.

## Author contributions

Conceptualization and designing the experiment: PBK and RV. Methodology: NASH, FG, RV and PBK. Data validation and analysis: NASH, FG, SG, RV and PBK. Investigation: NASH, PBK and FG. Resources: RV, RO, and PBK. Writing - original draft preparation: NASH and PBK. Writing - review and editing: NASH, RRV, SW, SG, RO, and PBK. Supervision and project administration: PBK, RV, SW, and RO. Funding acquisition: PBK, RV and RO. All authors contributed to the article and approved the submitted version.

## Acknowledgements

This research work was supported by FORMAS (2019-01316) and the Swedish Research Council (2019–04270), Novo Nordisk Fonden (0074727), Carl Tryggers Stiftelse (CTS 20:464) and Partneskap Alnarp (PA1365-2021). SCW acknowledges funding from the Scottish Government Rural and Environment Science and Analytical Services (RESAS) Division.

---

**Supplementary Table 1.** Analysis of soil nutrients and characteristics after amendment with selected rhizobacteria and growth of tomato and potato plants. TN: total nitrogen, P: phosphorus, K: potassium, EC: electrical conductivity, SOM: soil organic matter.

| Treatment | TN (mg/kg) |  | Available P (mg/l) |  | Available K (mg/l) |  | pH |  | EC (mS/cm) |  | SOM (%) |  |
| --- | --- | --- | --- | --- | --- | --- | --- | --- | --- | --- | --- | --- |
|  | Tomato | Potato | Tomato | Potato | Tomato | Potato | Tomato | Potato | Tomato | Potato | Tomato | Potato |
| *P. fluorescens* SLU99 | 7080 | 5780 | 36 | 29 | 140 | 15 | 5.9 | 6.5 | 1.7 | 1.4 | 42.7 | 43 |
| *S. rubidaea* EV23 | 6190 | 6000 | 34 | 27 | 140 | 11 | 5.9 | 6.4 | 2.1 | 1.3 | 43.4 | 48 |
| *S. rubidaea* AV10 | 6360 | 6030 | 34 | 27 | 130 | 14 | 5.9 | 6.4 | 2.4 | 1.4 | 39.6 | 44 |
| *S. plymuthica* S412 | 5510 | 6190 | 34 | 28 | 150 | 11 | 5.8 | 6.5 | 2.3 | 1.3 | 41.0 | 44 |
| Control | 6850 | 6160 | 32 | 28 | 130 | 9 | 5.9 | 6.5 | 2.3 | 1.2 | 42.9 | 46 |

**Supplementary Table 2.** Overview of differentially expressed transcription factor (TF) families identified in tomato and potato plants upon treatment with culture filtrate of *Pseudomonas fluorescens* SLU99.

| TF Family | Differentially Expressed TFs |  |  |  |
| --- | --- | --- | --- | --- |
|  | Tomato Root | Potato Root | Tomato leaf | Potato leaf |
| AP2 | 0 | 5 | 0 | 0 |
| ARR-B | 1 | 0 | 2 | 2 |
| B3 | 4 | 4 | 1 | 3 |
| bHLH | 11 | 23 | 19 | 8 |
| bZIP | 1 | 2 | 1 | 0 |
| CO-like | 0 | 6 | 1 | 3 |
| DBB | 0 | 5 | 5 | 0 |
| ERF | 16 | 18 | 15 | 4 |
| GATA | 2 | 6 | 5 | 3 |
| GRAS | 3 | 6 | 3 | 1 |
| HD-ZIP | 11 | 23 | 1 | 6 |
| HSF | 4 | 1 | 5 | 2 |
| LBD | 6 | 13 | 6 | 6 |
| MIKC_MADS | 14 | 8 | 1 | 2 |
| MYB | 21 | 16 | 16 | 8 |
| MYB_related | 3 | 8 | 2 | 5 |
| NAC | 2 | 12 | 3 | 8 |
| NF-Y | 4 | 4 | 3 | 1 |
| TALE | 2 | 4 | 1 | 1 |
| TCP | 0 | 5 | 0 | 2 |
| Trihelix | 1 | 1 | 1 | 1 |
| WOX | 1 | 1 | 0 | 0 |
| WRKY | 4 | 8 | 1 | 4 |
| ZF-HD | 1 | 3 | 0 | 0 |

**Supplementary Figure 1.**
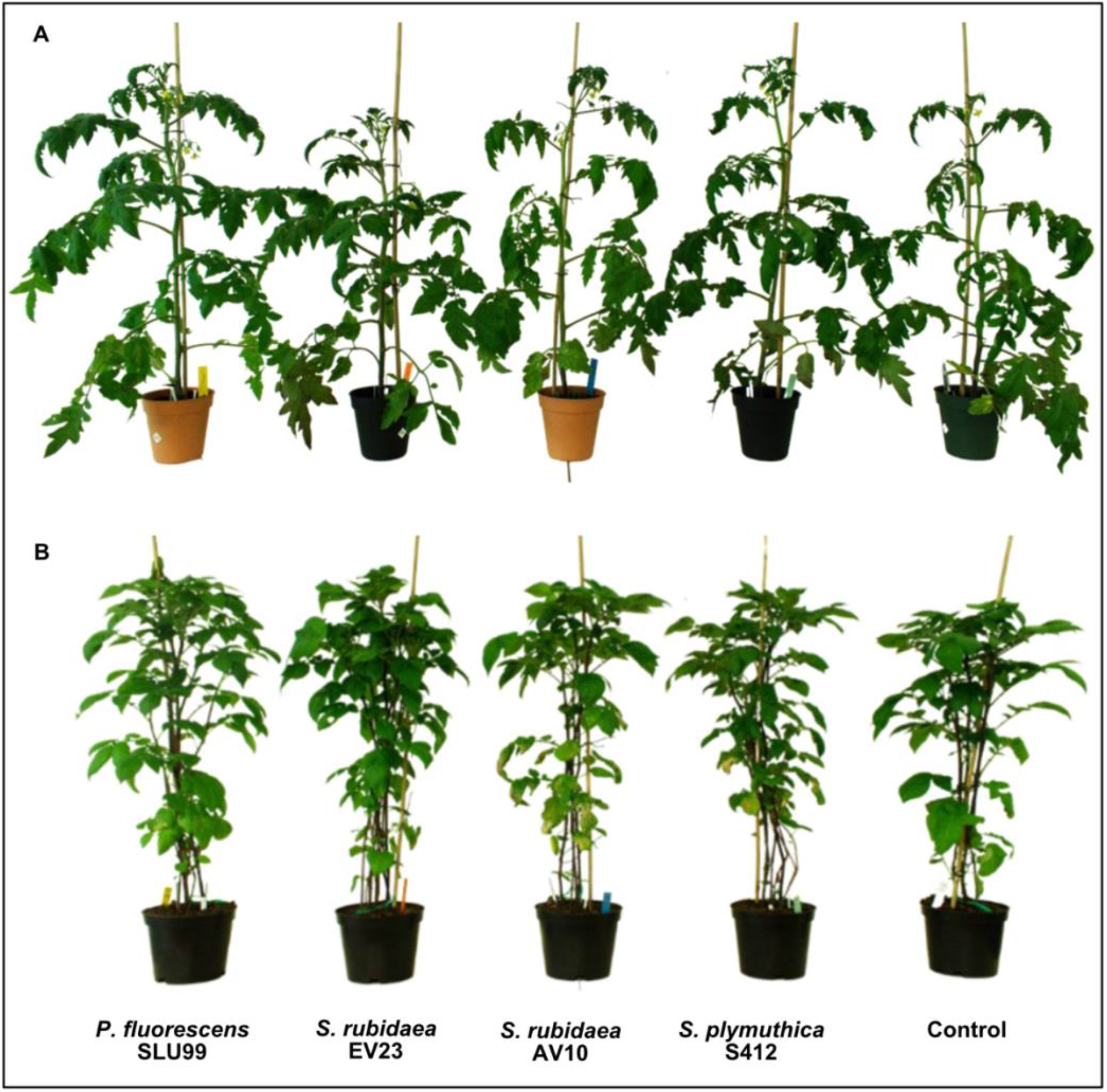
Plant height after treatment with different bacteria. (A) Tomato plants 42 days after transplanting (B) Potato plants 53 days after planting.

